## Supplementary figures and images for "Using deep mutational scanning to benchmark variant effect predictors and identify disease mutations"

### Fig EV1

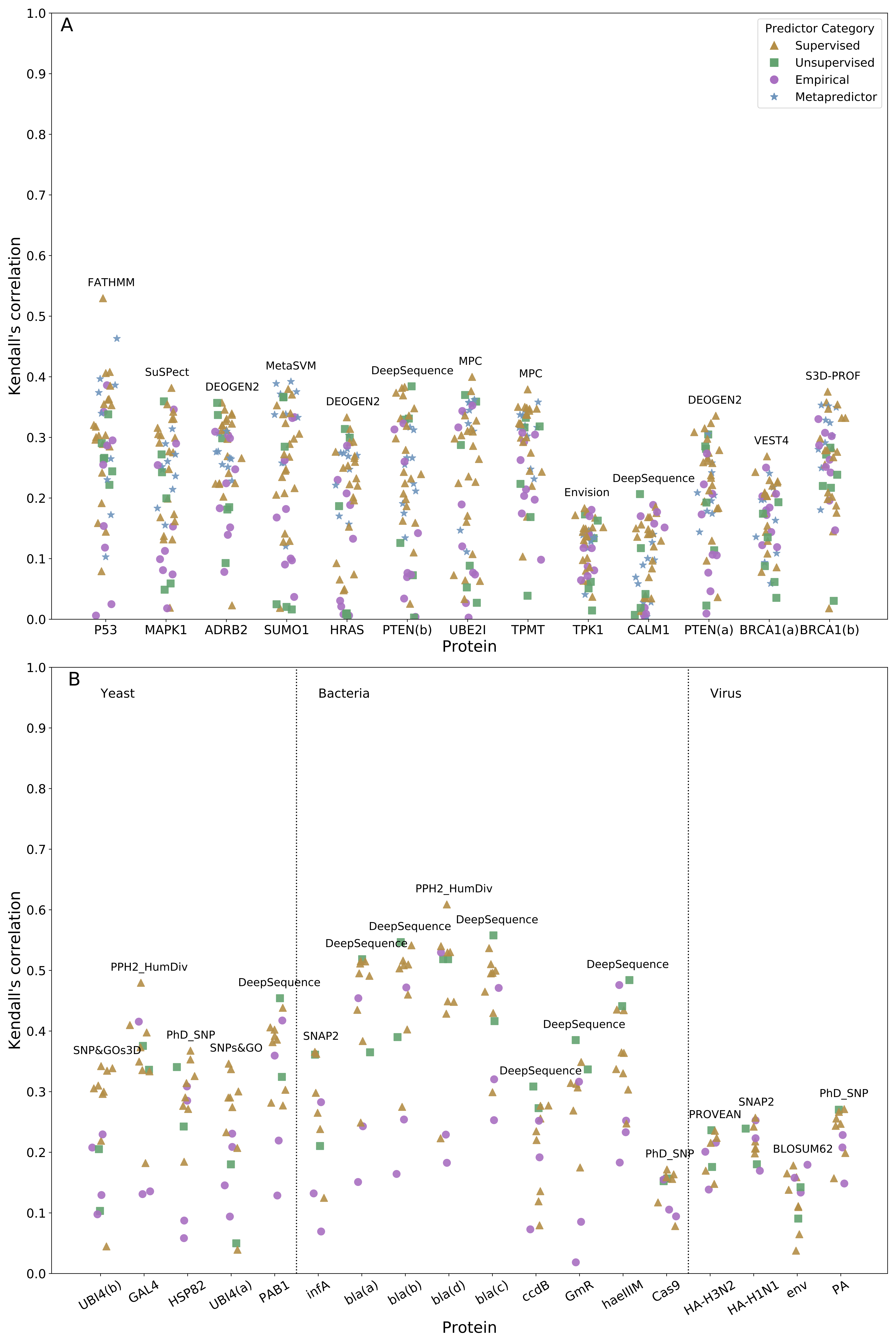

### Fig EV2

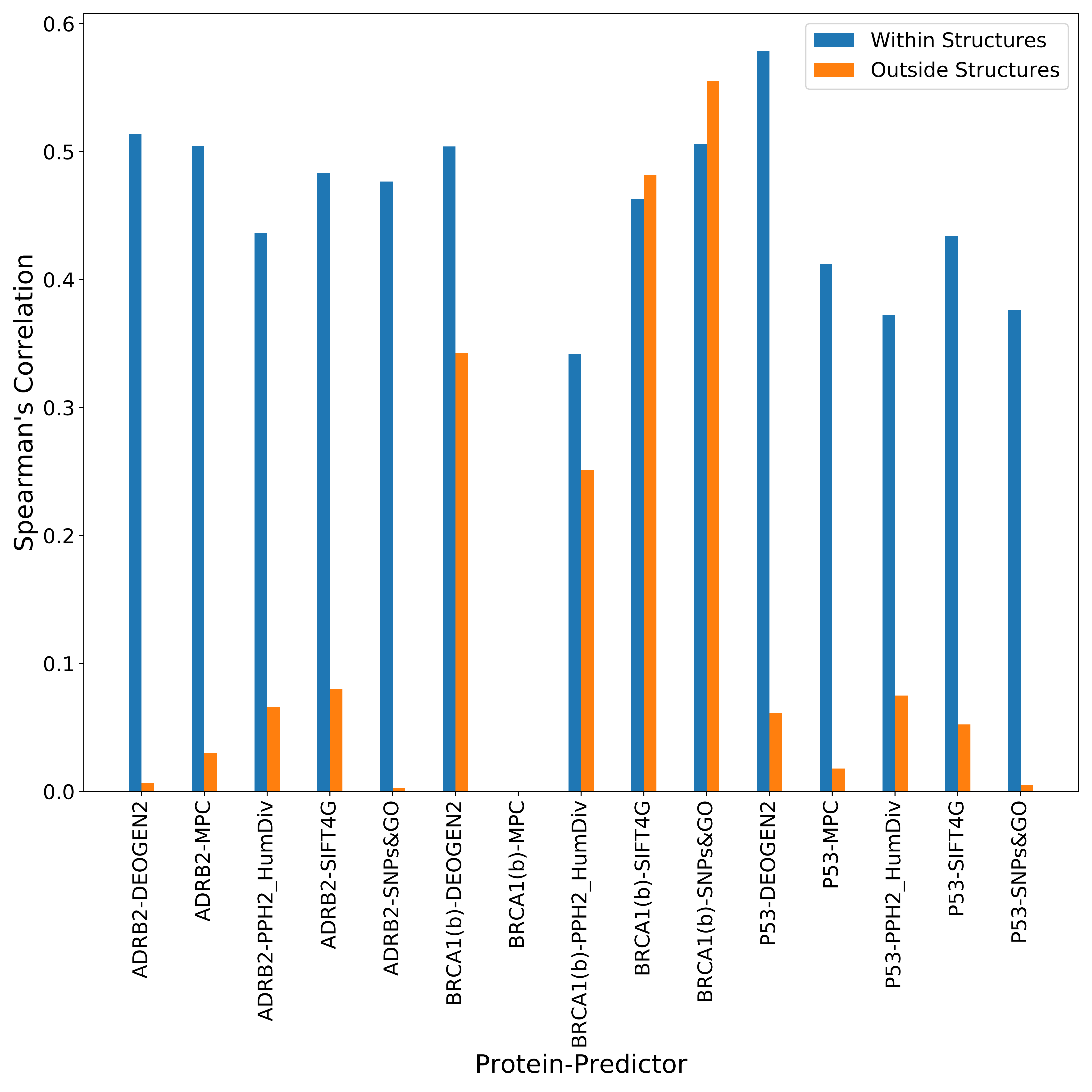

### Fig EV3

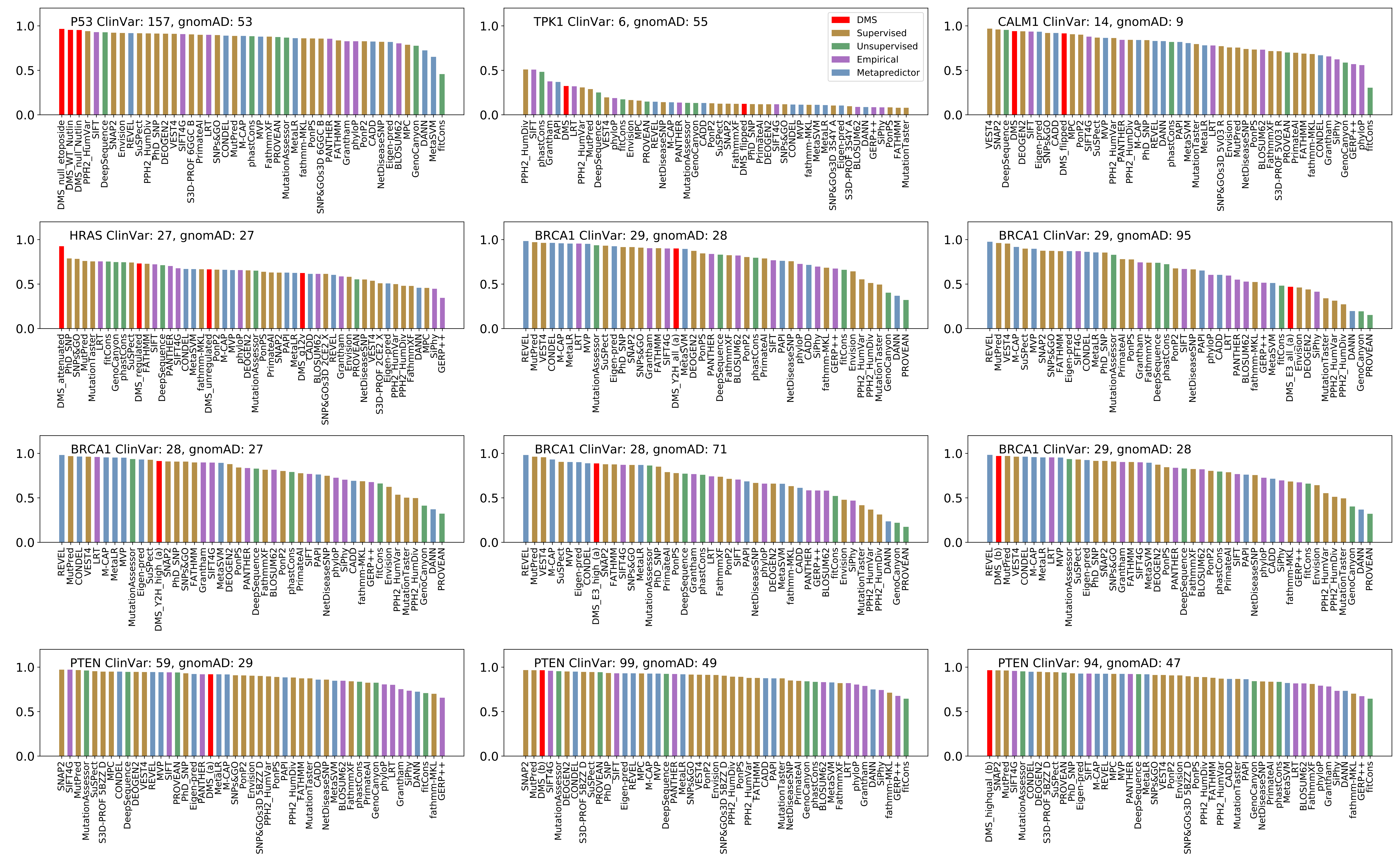

### Fig EV4

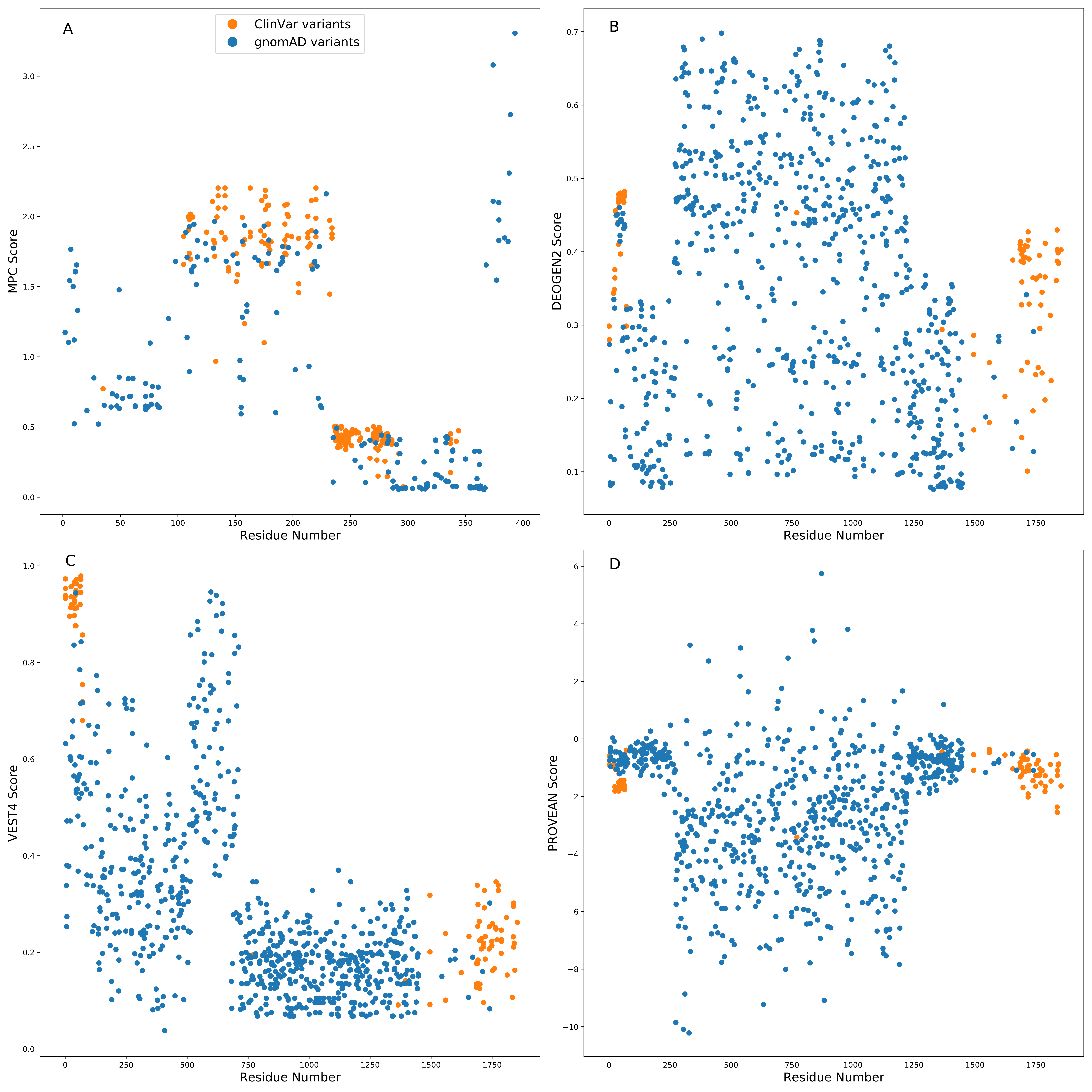
